# mTORC1 activates PASK-Wdr5 signaling to epigenetically connect the nutrient status with myogenesis

**DOI:** 10.1101/232173

**Authors:** Chintan K. Kikani, Xiaoying Wu, Sarah Fogarty, Seong Anthony Woo Kang, Noah Dephoure, Steve Gygi, David Sabatini, Jared Rutter

## Abstract

In the tissue microenvironment, stem cell functions are modulated by extrinsic signaling cues such as peptide hormones and dietary nutrients. These signaling cues maintain the balance between self-renewal and differentiation of its resident stem cells. The mechanistic Target of Rapamycin Complex 1 (mTORC1) is implicated to play an important role in regulating this balance, although its downstream effectors in stem cells have been elusive. We have recently shown that the PASK protein kinase phosphorylates Wdr5 to stimulate muscle stem cell differentiation by epigenetically activating the Myogenin promoter. Here, we show that the PASK-Wdr5 signaling pathway is a nutrient-sensitive downstream target of mTORC1 in muscle stem cells. We show that phosphorylation of PASK, and in turn of Wdr5, by mTORC1 is required for the activation of Myogenin transcription, exit from the self-renewal and induction of the myogenesis program. Thus, mTOR connects the diverse extrinsic signaling cues to a central epigenetic process to regulate the muscle stem cell fate between self-renewal and differentiation.

---

Skeletal muscle has a remarkable ability to restore its form and function following nearly complete myofiber destruction due to injury (Baghdadi and Tajbakhsh, 2017). This regenerative potential of skeletal muscle is largely attributed to its resident Muscle Stem Cells (MuSCs) (Chang and Rudnicki, 2014). MuSCs occupy a specific niche in the basal lamina, which supports their metabolic and cell cycle quiescence in uninjured muscle. Upon injury to myofibers, disruption of the niche triggers the activation of transcriptional, metabolic and signaling events within MuSCs resulting in cell division. The progeny of these proliferative cells ultimately undergo myogenic differentiation and fuse to regenerate the multi-nuclear myofibers (Buckingham and Rigby, 2014; Chang and Rudnicki, 2014; Relaix and Zammit, 2012).

Regenerative myogenesis is a well-coordinated program that involves the sequential action of multiple transcription factors working in concert with epigenetic regulators. Following an injury, quiescent Paired Box 7+ (Pax7+) MuSCs begin to proliferate, and a subset of these MuSCs gain expression of the bHLH transcription factor MyoD. Myogenin (MyoG) is a transcriptional target of MyoD and MyoD+/MyoG+ cells form differentiation-committed myoblasts and initiate the myoblast fusion program. Thus, induction of *Myog* expression is a key, irreversible step that establishes the myogenesis program. Thus, to ensure precise regulation of the *Myog* promoter activation, the epigenetic regulators such as histone methyl-transferases (HMTs), demethylases (KDMs), histone acetyltransferases (HATs), and deacetylases (HDACs) establish the framework for MyoD transcriptional function (Blum and Dynlacht, 2013; Saccone and Puri, 2010; Segales et al., 2014). In particular, histone H3 Lysine 4 methyl-transferase (H3K4me) activities of the Mixed Lineage Leukemia (MLL) enzymatic complexes are required for activation of the *Myog* locus during myogenesis (Aziz et al., 2010; Buckingham and Rigby, 2014; Rampalli et al., 2007). However, it remains incompletely understood how diverse niche-derived signaling cues impinge upon MLL complexes to regulate transcriptional activation of the *Myog* promoter.

Niche-derived signaling cues such as Wnt, Insulin, Insulin-like growth factors (IGFs) and nutrients are known to regulate MuSC activation, proliferation, commitment and finally the execution of the myogenesis program (Bentzinger et al., 2010; Dayanidhi and Lieber, 2014; Kuang et al., 2008). In particular, the establishment of myogenic commitment is regulated by the PI3K/Akt, mTOR, MAPK and β-catenin signaling pathways (Dayanidhi and Lieber, 2014; Dumont et al., 2015; Ge and Chen, 2012). The mTOR protein kinase, in particular, has been shown to be pivotal in MuSC function (Ge and Chen, 2012; Rodgers et al., 2014; von Maltzahn et al., 2011; Zhang et al., 2015). This kinase exists in two functionally distinct complexes, the Raptor-containing mTOR complex 1 (mTORC1) and the Rictor-containing mTOR complex 2 (mTORC2) (Laplante and Sabatini, 2012). mTORC1 is activated by nutrients such as amino acids and glucose, as well as Insulin/IGF-1 (Saxton and Sabatini, 2017) and Wnt signaling (Choo et al., 2006; Inoki et al., 2006), all of which are enriched in the MuSC niche. The loss of mTOR inhibits both MuSC proliferation and differentiation (Zhang et al., 2015), and this appears to be mostly explained by the loss of the Raptor-containing mTORC1 (Bentzinger et al., 2008). The genetic ablation of Rictor in MuSCs, however, appears to be well tolerated, although it may affect MuSC lineage specification (Hung et al., 2014). In addition to its function in regenerative myogenesis, recently mTORC1 was also implicated in inducing a G^alert^ state in MuSCs. G^alert^ is a quasi-activated state of MuSCs in an uninjured, contralateral leg in response to a muscle injury in a distinct leg (Rodgers et al., 2014). MuSCs in G^alert^ state show faster cycling times and increased *Myogenin* (*Myog*) expression. Thus, mTORC1 is critical for MuSC activation and regenerative myogenesis in response to injury. However, despite its importance, it remains unclear how mTORC1 signals to activate the myogenic transcriptional network.

We have recently identified a novel signaling pathway downstream of the PASK protein kinase, which connects signaling cues to phosphorylation of Wdr5, a member of MLL, SET1 and other histone-modifying enzymatic complexes, to drive transcriptional activation of *Myog* and myogenesis (Kikani et al., 2016). Our data show that PASK, via Wdr5 phosphorylation, collaborates with MyoD for transcriptional activation of *Myog* to drive the myogenesis program (Kikani et al., 2016). In response to differentiation cues, PASK phosphorylates Wdr5 at Ser49, which leads to enhanced MyoD recruitment and activation of the *Myog* promoter. Thus, we hypothesized that PASK-Wdr5 are intermediates of the signaling pathways that drive myogenesis (Kikani et al., 2016). However, it remained unclear how differentiation signaling cues might activate the PASK-Wdr5 pathway. Here, we identify PASK as an interacting partner and substrate of mTORC1 that is a necessary mediator of its myogenic function. Our data suggest that mTORC1 connects niche-derived nutrient sufficiency and hormonal signals to epigenetic complexes such as MLL via PASK phosphorylation to drive MuSCs differentiation.

## Results

Nutrients and insulin activate PASK in an mTORC1-dependent manner

We have previously reported that PASK expression was induced several-fold upon skeletal muscle injury and that loss of PASK resulted in severe defects in muscle regeneration (Kikani et al., 2016). We showed that PASK activity was also post-translationally stimulated during *in vitro* myogenesis (Kikani et al., 2016). To understand if this activation of PASK is part of its pro-myogenic function, we first asked if PASK is similarly activated during muscle regeneration *in vivo*. To do so, we generated mice expressing V5-tagged human PASK (hPASK) from the *Rosa26* locus (termed *Rosa26^hPASK-V5^*). Parenthetically, probably due to its post-translational regulation, over-expression of human PASK in mice did not result in any overt skeletal muscle phenotype in uninjured animals. However, during regeneration, *Rosa26^hPASK-V5^* mice showed a modest elevation of *Myog* mRNA and the MyoG target myosin heavy chain *(Myh3)* (Figure S1A) consistent with the known function of PASK to enhance myogenic gene expression. Consistent with the enhanced regenerative program in mice with PASK overexpression, MuSCs derived from the *Rosa26^hPASK-V5^* mice showed morphological evidence of differentiation as early as one day after isolation in culture media whereas wild-type MuSCs remained mononucleated for up to two days (Figure S1B). Using this PASK allele, which is not subject to transcriptional regulation (Figure S1A, compare hPASK vs. mPASK mRNA levels), we measured PASK kinase activity during tibialis anterior (TA) muscle regeneration. As shown in Figure 1A, PASK activity, as assessed by autophosphorylation (Kikani et al., 2010; Kikani et al., 2016), was induced 3 days after injury. The increase in PASK activity coincided with the time point when both MyoG and endogenous mouse PASK expression is induced (Figure 1A, Figure S1A).

**Figure 1.**
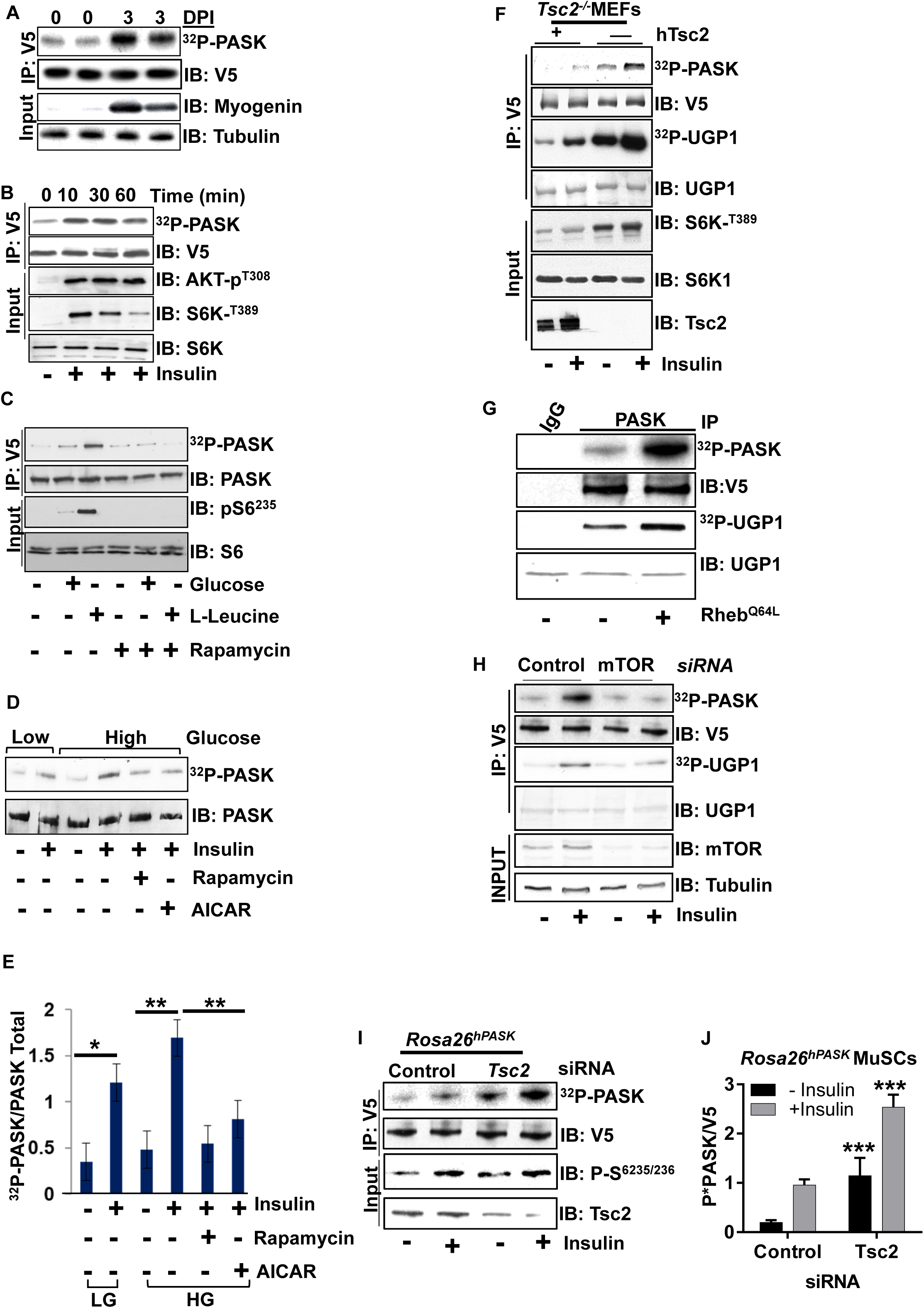
Nutrient and insulin signaling activate PASK via mTORCl. **[A]** PASK is activated during skeletal muscle regeneration. Left and right leg TA muscles were isolated from control or BaCh-injured *Rosa26^hPASK-V5^* mice 3 days post-injection. V5-tagged PASK was immunoprecipitated from tissue extract to assay PASK activation by an in vitro autophosphorylation assay which indicates the incorporation of ^32^P into PASK as a function of kinase activity. Immunoblot (IB) of MyoG marks myogenic regeneration. **[B]** CHO-K1 cells expressing V5-tagged human PASK were stimulated with 100nM insulin for the indicated times. PASK was immunoprecipitated using anti-V5 antibody and an *in vitro* kinase assay was performed as in [A]. Activation of PI-3K and mTORC1 signaling was demonstrated by appearance of phospho-AKT and phospho-S6K. **[C]** HEK293E cells were starved of amino acids and glucose for 8 hr, followed by stimulation with either 25 mM glucose or 800 μM L-leucine for 1 hr. Endogenous PASK was purified using anti-PASK antibody from cell extracts and *in vitro* kinase activity was performed as in [A]. **[D]** PASK from HEK293E cells was assayed as in [C]. Cells were stimulated with 100nM insulin for 1 hr after pretreatment with DMSO, 100nM Rapamycin or 25 μM AICAR for 4 hr. Endogenous PASK was purified using anti-PASK antibody from cell extracts and *in vitro* kinase activity assay was performed as in [A]. **[E]** Quantification of [D]. Phospho-PASK (^32^P-PASK) and total PASK were quantified from three independent experiments and the ratio expressed as the fold change in PASK activity under the indicated stimuli. Error bars are ± S.D. \**P*<0.05, \*\**P*<0.01. **[F]** V5-PASK was expressed in *Tsc2*^−/−^ cells with or without complementation with wild-type hTsc2. Cells were serum-starved overnight and then stimulated with 100nM insulin for 1hr. PASK was immunoprecipitated using anti-V5 antibody and *in vitro* kinase assay was performed. The yeast Ugp1protein, which is a robust *in vitro* substrate of PASK, was used as an exogenous substrate. **[G]** PASK was purified from HEK293T cells with a vector control (-) or expressing Rheb^Q64L^ and was purified and subjected to kinase activity assay as in **[F]**. **[H]** HEK293E cells were transfected with control or mTOR-targeting siRNA for 24 hr. A vector expressing V5-PASK was then transfected and, after 24 hr, cells were serum-starved overnight and then stimulated with 100nM insulin for 1 hr. PASK was purified and subjected to kinase activity assay as in **[F]**. **[I]** MuSCs isolated from *Rosa26^hPASK-V5^* mice were transfected with control or mouse Tsc2-targeting siRNA. 24 hr after transfection, cells were switched to 5% serum-containing medium overnight followed by 4 hr of total serum starvation. Cells were then treated with vehicle or 100nM insulin for 1 hr. PASK was then purified and subjected to kinase activity assay as in **[F]**. **[J]** Quantification of PASK kinase activity measurements from three experiments as in **[I]**. \**P*<0.05, \*\**P*<0.005.

To understand how niche signals, like nutrients and insulin, activate PASK during muscle regeneration, we examined PASK activation by these stimuli in a cell-autonomous manner. As shown in Figure 1B, PASK was acutely and transiently activated by insulin stimulation in CHO-K1 cells. Similarly, glucose and amino acids such as L-leucine, which are enriched in the stem cell niche, also activate PASK in HEK293E cells (Figure 1C). While glucose activated PASK modestly but consistently, we observed a strong increase in PASK activity upon addition of L-leucine to cell culture media (Figure 1C). Since mTOR complex 1 (mTORC1) is a convergence point in both insulin and amino acid signaling (Saxton and Sabatini, 2017), we asked if the mTORC1 activity is required for PASK activation. As shown in Figure 1C, the addition of the mTORC1 inhibitor rapamycin (Yip et al., 2010) completely blocked PASK activation by glucose and L-leucine.

To further explore the role of mTORC1 in the regulation of PASK activity downstream of nutrient and insulin stimulation, we analyzed PASK activity in the presence or absence of kinase modulators that either augment or inhibit mTORC1 activity. AMP-activated protein kinase (AMPK) is a negative regulator of mTORC1 kinase function (Inoki et al., 2012). As shown in Figure 1D-E, insulin and high glucose synergistically activated PASK, and this was suppressed by pretreatment with the AMPK activator 5-aminoimidazole-4-carboxamide ribonucleotide (AICAR) to an extent similar to rapamycin treatment. AMPK phosphorylates and activates Tsc2, which negatively regulates mTORC1 function (Inoki et al., 2003b). Consistent with that, *Tsc2*^−/−^ mouse embryonic fibroblasts (MEFs), complemented with empty vector control but not with human Tsc2 (Onda et al., 1999), showed mTORC1 hyperactivation as evidenced by increased phosphorylation of the mTORC1 substrate p70S6K (Figure 1F). Loss of *Tsc2* also increased PASK kinase activity as shown by increased *in vitro* autophosphorylation and phosphorylation of its heterologous substrate Ugp1 (Figure 1F) (Smith and Rutter, 2007). Tsc2 functions as a GTPase-activating protein (GAP) for the Rheb GTPase, which stimulates mTORC1 activity (Inoki et al., 2003a). Expression of a constitutively activated Rheb (Rheb^Q64L^), which hyperactivates mTORC1, also resulted in increased PASK activity (Figure 1G). On the other hand, silencing mTOR resulted in a near complete block of insulin-stimulated PASK activation (Figure 1H).

Finally, to test whether mTORC1 contributes to PASK activation in muscle stem cells, we isolated MuSCs from *Rosa26^hPASK-V5^* mice and assessed PASK activation by insulin after silencing *Tsc2.* Not surprisingly based on our previous results from injury experiments, PASK was activated during MuSC isolation but was further activated by insulin stimulation (Figure 1I, J). The loss of *Tsc2* resulted in PASK activation both in the presence or absence of insulin. Thus, our results demonstrate that the mTORC1 complex activates PASK in response to insulin and nutrient signaling.

### PASK is phosphorylated by mTORCl at multiple residues to stimulate its activity

We hypothesized that mTORC1-dependent phosphorylation of one or more residues on PASK might result in its activation by nutrients and insulin. Therefore, we performed metabolic in-cell labeling using radioactive (^32^P) phosphate in *Tsc2*^−/−^, or *hTsc2* complemented *Tsc2*^−/−^ MEFs in the presence or absence of rapamycin. As shown in Figure 2A, endogenous PASK derived from *Tsc2*^−/−^ MEFs showed significantly increased phosphorylation compared with h*Tsc2*-complemented cells. Furthermore, this increase in PASK phosphorylation was partially suppressible by low-dose rapamycin, consistent with some other mTORC1 substrates (Figure 2A) (Kang et al., 2013). We also tested if, similar to PASK activity (Figure 1G), PASK phosphorylation was also induced by constitutively-activated Rheb (Rheb^Q64L^) and if that is dependent upon the catalytic activity of PASK. As shown in Figure 2B, both WT and KD (kinase dead) PASK showed enhanced in-cell phosphorylation in the presence of Rheb^Q64L^. In contrast, the phosphorylation of PDK1, an upstream activator of Akt that is not an mTORC1 substrate, was not induced by Rheb co-expression. Thus, mTORC1 activation induces the phosphorylation of PASK in cells, and the increased PASK phosphorylation is independent of its own catalytic activity, demonstrating that autophosphorylation is not required.

**Figure 2.**
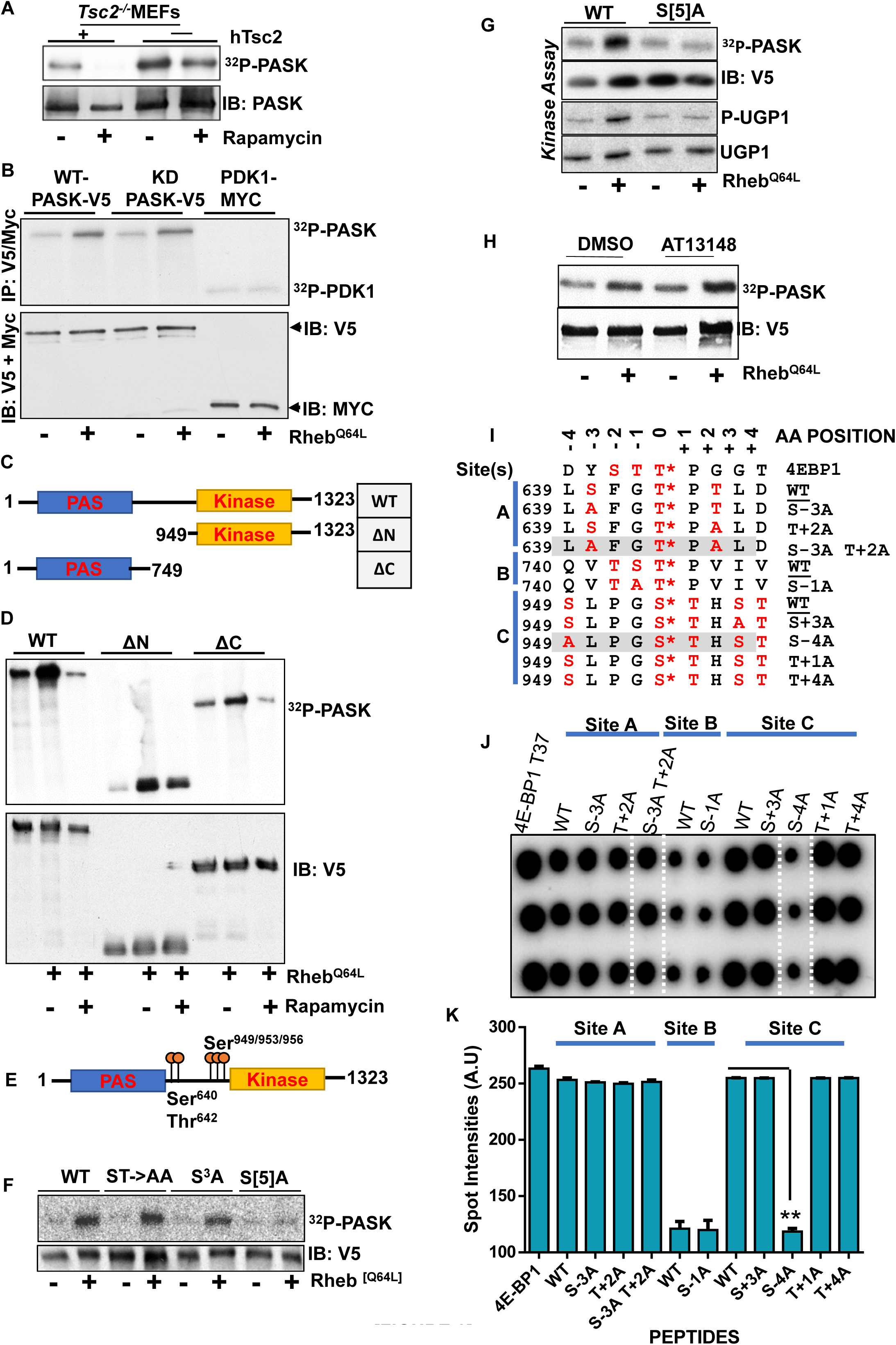
PASK is a direct phosphorylation target of mTORC1. **[A]** Endogenous mouse PASK was immunoprecipitated from *Tsc2*^−/−^ or *hTsc2-* complemented *Tsc2-^/^-* MEFs after labeling with ^32^P-phosphate for 4hr. Anti-PASK immunoprecipitates were separated by SDS-PAGE and subjected to autoradiography and immunoblot (IB). **[B]** HEK293T cells were transfected with vectors expressing V5-tagged WT or K1028R (kinase dead; KD) PASK or Myc-tagged PDK1 as well as either empty vector or a vector expressing Rheb^Q64L^. At 24 hr post-transfection, in cell ^32^P labeling was conducted and the indicated immunoprecipitates were analyzed as in **[A]**. **[C]** Schematic indicating the domain structure of full-length PASK and the domain truncation mutants used in **[D]**. **[D]** WT PASK or the truncation mutants from **[C]** were co-expressed with empty vector or a vector expressing Rheb^Q64L^ and were treated with or without 40 nM rapamycin. They were assessed for in cell phosphorylation as in **[A]**. **[E]** Schematic of the mTORC1-dependent phosphorylation sites on PASK. [F] WT or ST->AA (S^640^A T^642^A), S3A (S^949^A S^953^A S^956^A) or S[5]A (S^640^, T^642^, S^949^, S^953^, S^956^ to Ala) mutants of PASK were expressed in HEK293T cells with or without expression of Rheb^Q64L^ and analyzed for in cell phosphorylation as in **[A]**. **[G]** WT or the S[5]A mutant of PASK were expressed in HEK293T cells with or without co-expression of Rheb^Q64L^ and kinase activity was measured by autophosphorylation (^32^P-PASK) and Ugp1 phosphorylation as in Figure 1F. **[H]** Rheb^Q64L^-induced PASK *in vivo* phosphorylation was measured with or without 50 μM AT13148 pre-treatment in HEK293T cells. **[I]** Depiction of three separate peptides (Site A, B, and C) analyzed for *in vitro* phosphorylation by mTORC1. Phosphorylatable residues or their mutant version (in red) are marked within each peptide. The position matrix from -4 to 4 indicates each residue that are mutated with respect to the central phospho-acceptor residue (position 0 indicated by *) in 4E-BP1 (T^37^). The gray highlight indicates peptides that identified critical phosphorylated residues. **[J]** Phosphorylation of peptide by mTORC1 as analyzed by dot-blot as described in methods. White vertical lines correspond to gray highlighted peptides in [I]. **[K]** Quantification of phosphorylation intensities of mutated peptide over its WT version from experiments in [J]; *P*<0.005.

To identify residues on PASK that are specifically targeted by mTORC1 activity, we performed a domain truncation analysis in the presence or absence of Rheb^Q64L^ and rapamycin (Figure 2C). Rheb stimulated PASK phosphorylation within both the C-terminal kinase domain-containing region including residues 949-1323 as well as the N-terminal fragment containing the first 749 residues (Figure 2D). Surprisingly, these two regions showed differences in sensitivity to rapamycin inhibition as the AC fragment showed much more rapamycin sensitivity than the ΔN fragment (see discussion). Using mass spectrometry, bioinformatics, and site-directed mutagenesis, we identified two clusters of sites that were hyperphosphorylated upon mTORC1 activation. The sites within the N-terminal fragment include Ser^640^ and Thr^642^ (Figure 2E). The C-terminal phosphorylation sites include Ser^949^, Ser^953^, and Ser^956^. Mutation to alanine of either cluster alone had minimal effect on PASK phosphorylation induced by Rheb^Q64L^ expression (Figure 2F). Mutation of all five sites (S^640^, T^642^, S^949^, S^953^, S^956^ to Ala, termed S[5]A) however, resulted in essentially complete inhibition of Rheb-stimulated PASK phosphorylation (Figure 2F) and kinase activation (Figure 2G).

mTORC1 activates multiple kinases within the AGC family of protein kinases, such as Akt, p70S6K, and p90RSK. However, inhibition of these AGC kinases with a pan-AGC kinase inhibitor AT13148 did not affect Rheb-stimulated PASK phosphorylation (Figure 2H). Interestingly, the sequence surrounding the Rheb-stimulated PASK phosphorylation sites appears similar to many of the recently identified mTORC1 substrates (Kang et al., 2013). To test if mTORC1 can directly phosphorylate these sites, we utilized a mutagenized peptide array system to identify novel mTORC1 substrates as previously described (Kang et al., 2013). The mutated peptide library was generated by mutating phosphorylable residues within each peptide except the phosphorylatable residue at position 0 (indicated by * in Figure 2I). These peptides were then used as substrates for *in vitro* kinase reactions with purified mTORC1. As shown in Figure 2J-K, a peptide in which Thr^642^ was the only phosphorylatable residue (in Site A sequence) showed similar phosphorylation to the WT peptide. Site B was used as a negative control, being a poor mTORC1 substrate in our experiments. On the other hand, Site C showed robust phosphorylation by mTORC1 and mutation of Ser^949^ was sufficient to significantly diminish mTORC1-mediated phosphorylation of this peptide (Figure 2J-K) *in vitro.* Taken together, our data suggest that mTORC1-mediated phosphorylation of PASK at multiple residues is required for nutrient and insulin signaling to activate PASK.

### PASK forms a nutrient-sensitive complex with mTOR complex 1

To understand the mechanistic basis whereby PASK is phosphorylated and activated by mTORC1, we sought to understand whether PASK associates with mTOR or any of its associated Complex 1 proteins. We first immunoprecipitated V5-tagged WT PASK or a K1028R mutant lacking kinase activity (KD) from cells that co-express either Myc-tagged WT mTOR or a D2357E/V2364I mutant lacking kinase activity (KD). Both WT and KD PASK could be immunoprecipitated with either the WT or KD version of mTOR (Figure 3A). Interestingly, the WT PASK associated with KD mTOR showed significantly diminished phosphorylation at Thr^307^, which is an autophosphorylation site that we have shown previously to be a reliable marker of PASK activity (Wu et al., 2014). Thus, KD mTOR appears to function as a dominant negative towards PASK activity. When endogenous mTOR was isolated from cells, PASK was co-purified in addition to the members of the mTORC1 complex (Figure 3B). Similarly, immunoprecipitation of PASK-V5 co-purified endogenous mTOR and raptor (Figure 3C). When mTOR was silenced, the association of PASK with mTOR and with Raptor was significantly reduced suggesting that the PASK-Raptor association is likely mediated by mTOR. Using domain truncation analysis, we found that the C-terminal residues 949-1323 in PASK, which include the kinase domain and surrounding regions, are necessary to interact with endogenous mTORC1 (Figure 3D). Finally, we found that a mixture of L-leucine and Larginine, as well as expression of Rheb^Q64L^, two of the stimuli that led to phosphorylation and activation of PASK, both weaken the PASK-mTOR association (Figure 3E). Thus, PASK appears to form a nutrient and signaling-sensitive complex with mTORC1, similar to what was previously reported for the mTOR-Raptor association (Kim et al., 2002). These data suggest that PASK dynamically associates with mTORC1 wherein mTOR directly phosphorylates and activates PASK, resulting in its release from the complex.

**Figure 3.**
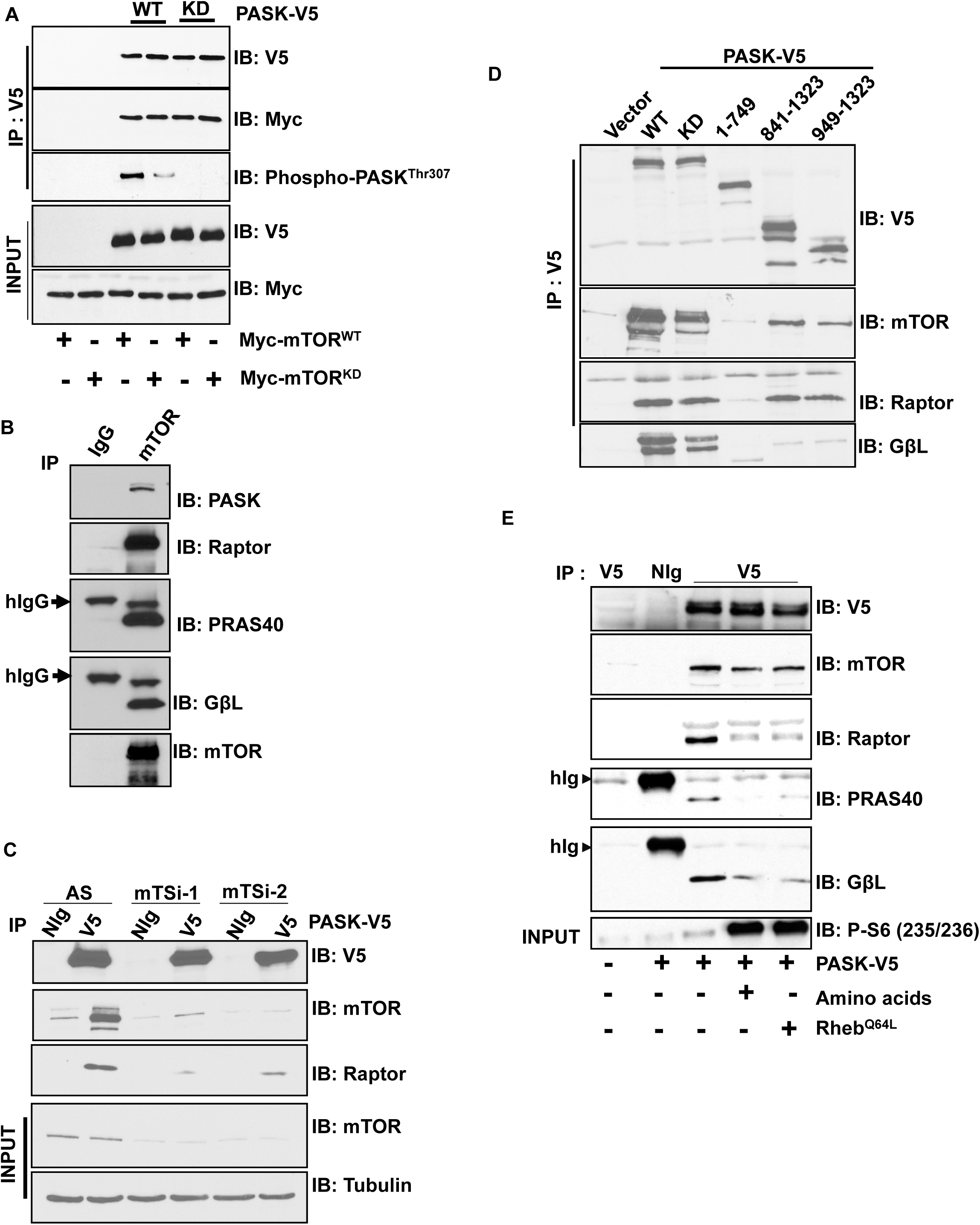
PASK associates with mTORCl in a nutrient-sensitive manner. **[A]** Vector control or WT or KD (K1028R) PASK was co-expressed with either Myc-tagged WT or D2357E (KD) mTOR in HEK293T cells. 24 hr after transfection, V5-tagged proteins were purified, and the presence of Myc-tagged mTOR was detected by western blotting. Immunoprecipitates were also probed with anti-AKT substrate antibody as described in methods to detect PASK-T307 phosphorylation. **[B]** mTOR protein was purified from HEK293T cells and the presence of its associated proteins was detected in the immunoprecipitates by western blotting. **[C]** mTOR silencing and V5-hPASK expression and immunoprecipitation were performed as described in Figure 1H. The presence of mTOR and its complex members were detected by western blotting of the immunoprecipitates. **[D]** The indicated V5-tagged PASK truncation mutants were expressed and immunoprecipitated from HEK293T cells. The co-IP of mTORC1 was determined by western blotting of the immunoprecipitates. **[E]** Vector or V5-PASK were expressed with Rheb^Q64L^ as indicated. For amino acid stimulation, cells were starved of amino acids L-leucine and L-arginine overnight. On the next day, 800 μM L-leucine and 100 μM L-arginine were added for 6 hr. Cells were lysed and V5-tagged PASK was purified from HEK293T cells. The relative abundance of mTORC1 was detected by western blotting of the immunoprecipitates.

### mTORCl and PASK activate the myogenesis program

*Myog* expression in myoblasts marks irreversible commitment to differentiate (Olguin et al., 2007). Hence, activation of *Myog* transcription is a major focal point of the signaling pathways that regulate myogenesis (Mok and Sweetman, 2011). As a tissue with significant metabolic demand, skeletal muscle homeostasis is tightly linked with nutrient status. This is consistent with the fact that the nutrient-responsive mTORC1 has been shown to regulate myogenesis and myofiber hypertrophy (Erbay and Chen, 2001; Ge and Chen, 2012; Saxton and Sabatini, 2017; von Maltzahn et al., 2011; Zhang et al., 2015) although its downstream effectors remain unknown (Ge and Chen, 2012). Because mTORC1 activates PASK and we previously demonstrated that PASK is required for efficient damage-induced myogenesis (Kikani et al., 2016), we hypothesized that mTORC1-dependent PASK activation is a mechanism whereby nutrient signaling could be coupled to myogenesis. We first compared myogenic induction upon loss of PASK and mTORC1 signaling in both MuSCs and C2C12 myoblasts. As shown in Figure 4A-B, both mTORC1 and PASK inhibition (with rapamycin and BioE-1197, respectively) effectively and similarly suppressed MyoG^+^ conversion and myoblast fusion in isolated MuSCs. This failure to convert to MyoG^+^ cells appears to be due to impaired induction of *Myog* mRNA in the presence of the inhibitors (Figure 4C). We next sought to identify components of the mTORC1 and mTORC2 complexes that are necessary for MyoG expression and myogenesis. To do so, we silenced mTOR, Raptor (member of mTORC1), Rictor (member of mTORC2) or the mTORC1 substrates, p70S6K (major substrate of mTORC1) or PASK during insulin-stimulated myogenesis. Consistent with a previous report (Bentzinger et al., 2008), mTORC1 but not mTORC2 is required for MyoG protein expression as loss of Raptor but not Rictor suppressed MyoG induction (Figure 4D). Furthermore, silencing of PASK, but not p70S6K, suppressed MyoG expression, suggesting that mTORC1-PASK but not mTORC1-S6K signaling is required for induction of the myogenic program.

**Figure 4.**
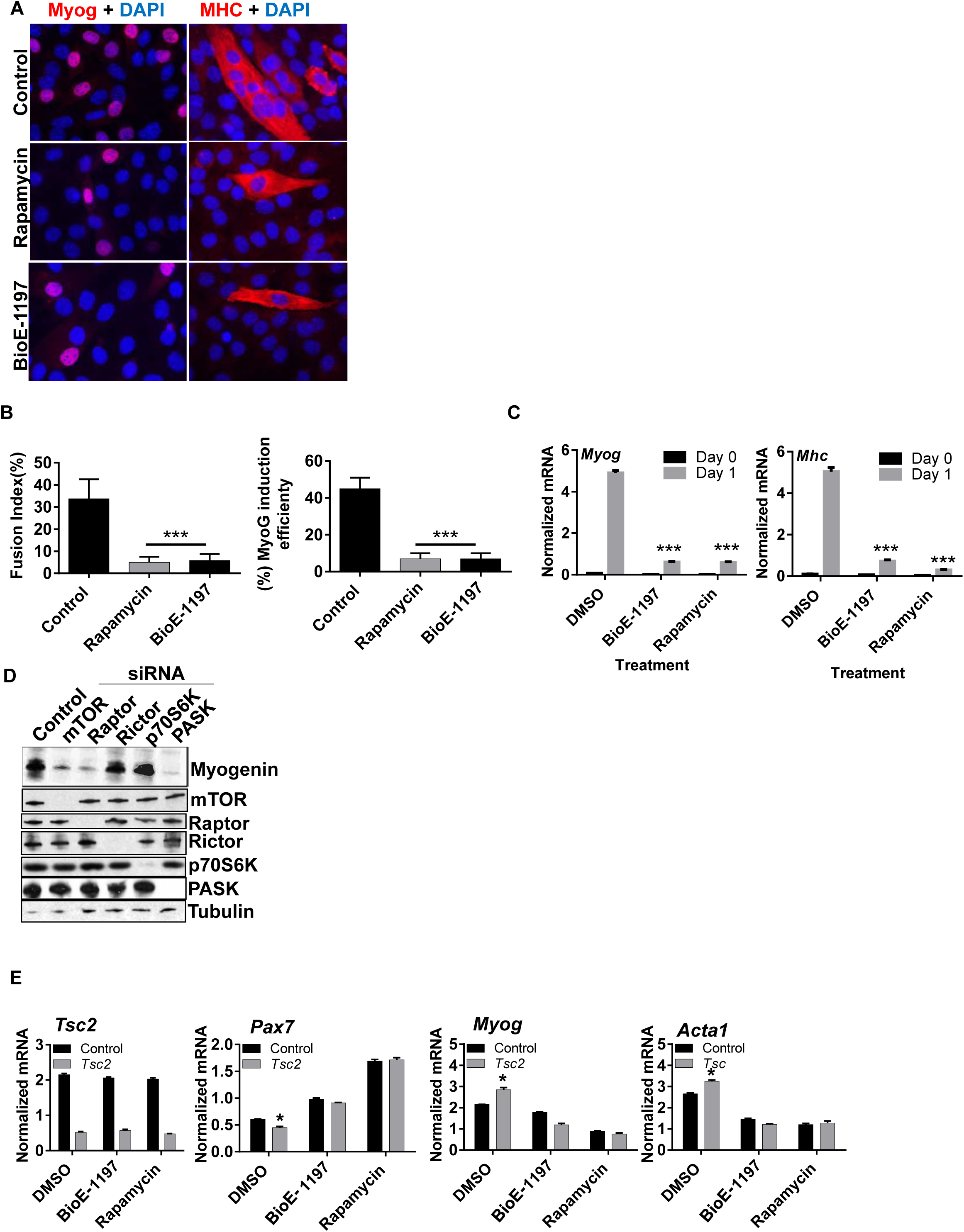
mTORC1 and PASK are required for efficient myogenesis. **[A]** MuSCs were isolated from TA muscles of WT mice and 24 hr after isolation, cells were treated with 100 nM rapamycin or 50 μM BioE-1197. 24hr after inhibitor treatment, cells were induced to differentiate by adding 100 nM Insulin. Cells were fixed after 1 day of differentiation and stained using anti-Myogenin or anti-MHC antibody for immunofluorescence. **[B]** Fusion index was calculated from experiments as in [A]. The fusion index is defined as the fraction of total nuclei that are found within myotubes. Similarly, Myogenin induction efficiency was determined by calculating the number of MyoG+ nuclei vs. total nuclei in the population. **[C]** mRNA was prepared from an experiment as in [A] and levels of *Myog* and *Mhc (Mylpf)* were measured using qRT-PCR and normalized using 18s rRNA. **[D]** The indicated components of mTORC1 and mTORC2 were silenced in C2C12 myoblasts. 48hr after silencing, cells were stimulated to differentiate using 100nM insulin. Myogenin expression was used as an indication of differentiation for each cell population. **[E]** *Tsc2*-targeted or control siRNA was transfected into C2C12 cells. 24 hr after transfection, cells were treated with 100 nM rapamycin or 50 μM BioE-1197. 24 hr after drug treatment, cells were induced to differentiate using 100 nM insulin in the continued presence of inhibitors. Normalized mRNA levels of indicated transcripts from these cells were measured by qRT-PCR.

Finally, to determine how mTORC1 activation affects myogenesis and what role PASK plays in that process, we silenced *Tsc2* in cultured myoblasts and analyzed the mRNA levels for *Pax7, Myog,* and *Acta1* which marks proliferating, committed and differentiated stages, respectively. As shown in Figure 4E, loss of *Tsc2* resulted in a modest but significant decrease in *Pax7* mRNA and an increase in *Myog* mRNA, suggesting mTORC1 activation increases myogenic commitment. This is further evidenced by increased expression of the muscle-specific actin *Acta*, which marks terminally differentiated myocytes. This effect of mTORC1 hyperactivation on increased commitment and myogenesis was completely suppressed by either inhibition of PASK or mTORC1. These data suggest that mTORC1 and PASK are both required for myogenin expression during myogenesis and that PASK might be a novel mTORC1 substrate and effector in this process.

### PASK is a downstream target of mTORCl to stimulate myogenic initiation

MuSCs from *Rosa26^hPASK-V5^* mice show enhanced myogenesis compared with control MuSCs as indicated by increased MyoG staining (Figure 5A-B) and fusion index (% of nuclei inside myotubes/total number of nuclei, Figure 5C-D). Rapamycin treatment effectively reversed this increase in myogenesis in PASK over-expressing MuSCs. We reasoned that since rapamycin is able to suppress myogenesis in MuSCs from *Rosa26^hPASK-V5^* mice, mTORC1 is likely already activated during isolation of MuSCs. If so, *Tsc2* silencing should not have an additive effect on myogenesis, as PASK is already activated by mTORC1. Consistent with that, while the activation of the mTORC1 pathway by *Tsc2* silencing resulted in modest stimulation of myogenesis in control cells (Figure 5A-D), MuSCs from *Rosa26^hPASK-V5^* did not show further enhancement of myogenesis. Again, rapamycin treatment inhibited myogenesis regardless of PASK over-expression, suggesting the requirement for activated mTORC1 in inducing myogenesis downstream of PASK.

**Figure 5.**
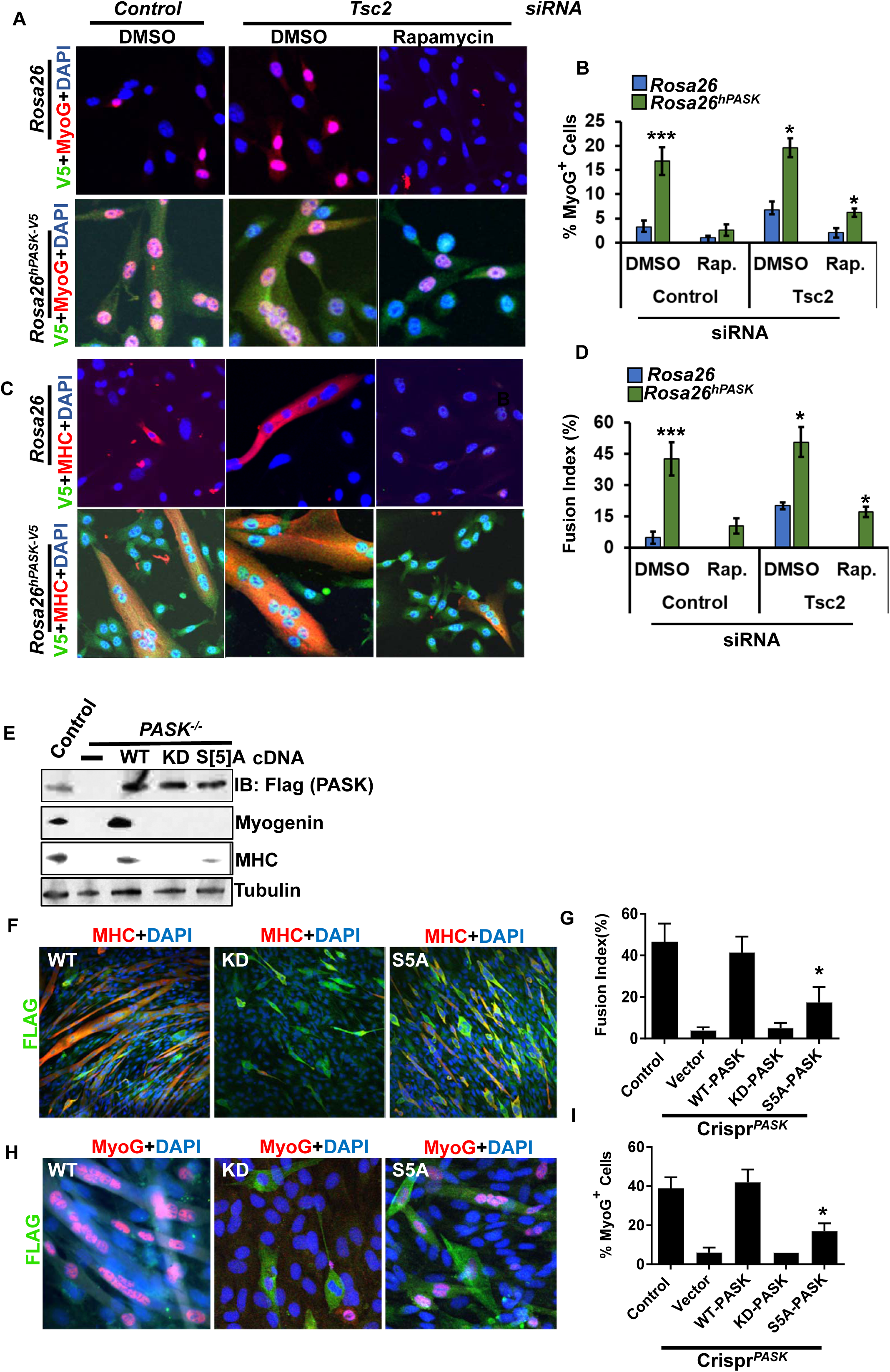
PASK phosphorylation at mTORC1 sites is required for efficient myogenesis. **[A-D]** MuSCs isolated from *Rosa26^hPASK-V5^* mice were transfected with control or *Tsc2-* targeting siRNA. 24 hr after transfection, cells were treated with either DMSO or with rapamycin in growth media. 48 hr later, cells were fixed and analyzed for **[A-B]** levels of MyoG and **[C-D]** the extent of myogenesis by measuring the fusion index. **[E]** PASK^+/+^ or *PASK*^−/−^ MuSCs were isolated from TA muscles of mice with the respective genotype. 24 hr after isolation, *PASK*^−/−^ cells were infected with retroviruses expressing Flag tagged WT, K1028R or S[5]A (S^640^, T^642^, S^949^, S^953^, S^956^ to Ala) PASK. 48 hr after infection, PASK+^/^+ and *PASK*^−/−^ cells were stimulated to differentiate with 100 nM insulin. Protein extracts were prepared from all cells and myogenesis was measured by western blotting using the indicated antibodies. **[F]** C2C12 myoblasts with deletion of PASK using CRISPR/Cas9 were infected with the indicated retroviruses. 48 hr after retroviral infection, C2C12 cells were induced to differentiate using 100 nM Insulin. Myogenesis was determined by immunofluorescence microscopy using antibodies against MHC. **[G]** Fusion index was calculated as in Figure 4B. **[H-I]** Cells from an experiment as in [F] were stained with anti-MyoG antibody to determine MyoG induction efficiency as described in Figure 4[B].

To determine if PASK phosphorylation by mTORC1 is required for efficient myogenesis, we retrovirally expressed vector control, WT, KD or S[5]A-mutated PASK in isolated MuSCs from *PASK*^−/−^ mice. As shown in Figure 5E, the re-expression of WT, but not KD, PASK fully restored MyoG and MHC expression in isolated *PASK*^−/−^ MuSCs. Expression of S[5]A-mutated PASK, on the other hand, had very modest effects on MyoG and MHC (Figure 5E). We also utilized the CRISPR/Cas9 system to delete endogenous mouse PASK in C2C12 myoblasts (Crispr^*PASK*^) and reconstituted with either empty vector or WT, KD or S[5]A-mutated PASK. Using these cell lines, we assayed the effectiveness of myogenesis in response to insulin. Expression of WT, but not KD, PASK resulted in the full restoration of myogenesis as measured by the fusion index (Figure 5F-G). Again, S[5]A-mutated PASK only modestly rescued the defect in myotube formation caused by loss of PASK. Similar results were obtained for MyoG expression, where WT PASK fully rescued the defect in MyoG expression and PASK lacking the mTORC1 phosphorylation sites was impaired in that function (Figure H-I). Thus, PASK and specifically the residues phosphorylated by mTORC1 are required for the efficient induction of myogenesis by insulin.

### PASK phosphorylation of Wdr5 is required for mTORC1-stimulated myogenesis

We have previously shown that phosphorylation of Wdr5 is necessary and sufficient for the effects of PASK to induce *Myog* transcription and myogenesis (Kikani et al., 2016). Moreover, the physical interaction between PASK and Wdr5 was specifically induced upon insulin treatment to initiate myogenesis (Kikani et al., 2016). Herein, we showed that insulin treatment also stimulated PASK phosphorylation and activation in an mTORC1-dependent manner. Hence, we hypothesized that mTORC1 activation might enhance the PASK-Wdr5 association. To test this, we co-expressed WT or KD PASK with WT Wdr5 in the presence or absence of Rheb^Q64L^ and measured the PASK-Wdr5 association. As expected, expression of Rheb^Q64L^ caused increased PASK activity as measured by autophosphorylation at Thr^307^ (Figure 6A). This was coincident with augmentation of the PASK-Wdr5 interaction. Interestingly, the C-terminal residues where mTORC1 phosphorylates PASK are adjacent to the Wdr5 binding region in PASK that we identified previously (Kikani et al., 2016) (Figure 6B). To more precisely map the Wdr5 binding region in PASK, we mutagenized the conserved residues within this stretch and examined effects on the interaction with Wdr5 (Figure 6C). As shown in Figure 6D, mutation of the highly conserved C^924^ and W^926^ PASK residues to alanine resulted in a significantly weakened interaction with Wdr5. As these residues are adjacent to the mTORC1 phosphorylation site on PASK (Figure 6C), we hypothesized that mTORC1-mediated PASK phosphorylation might augment Wdr5 binding. Indeed, we found that the S[5]A mutant, lacking mTORC1-mediated phosphorylation, failed to show Rheb^Q64L^-dependent induction of the PASK-Wdr5 interaction (Figure 6E). Taken together, these data suggest that mTORC1-mediated phosphorylation of PASK stimulates the PASK-Wdr5 association. We have shown previously that this interaction correlates strongly with PASK phosphorylation of Wdr5 at Ser^49^, which orchestrates epigenetic changes at the *Myog* promoter to enable gene expression. To test if Wdr5 phosphorylation at Ser^49^ by PASK is a mechanism whereby mTORC1 signals to induce myogenesis, we first tested whether expression of the phospho-mimetic Wdr5 mutant (Wdr5^S49E^) might rescue the defect in myogenesis resulting from mTORC1 inhibition. As shown in Figure 6F, rapamycin completely prevented the induction of *Myog* and *Mylff* in response to differentiation cues. These defects were completely reversed by expression of the phospho-mimetic Wdr5 mutant Wdr5^S49E^, while expression of Wdr5^WT^ or Wdr5^S49A^ had no effect (Figure 6F). Consistent with the mRNA data, western blot analysis showed that Wdr5^S49E^, but not Wdr5^S49A^ also restored MyoG and MHC protein induction in rapamycin-treated cells (Figure 6G). Finally, expression of Wdr5^S49E^, but not Wdr5^WT^ or Wdr5^S49A^, rescued the rapamycin-induced defect in myogenesis as assessed by formation of multicellular myotubes (Figure 6H-I). Taken together our results have identified a signaling pathway that transmits nutrient and hormonal signals via mTORC1 phosphorylation and activation of PASK to induce *Myog* expression and myogenic commitment through phosphorylation of the Wdr5 epigenetic regulator (Figure *7*).

**Figure 6.**
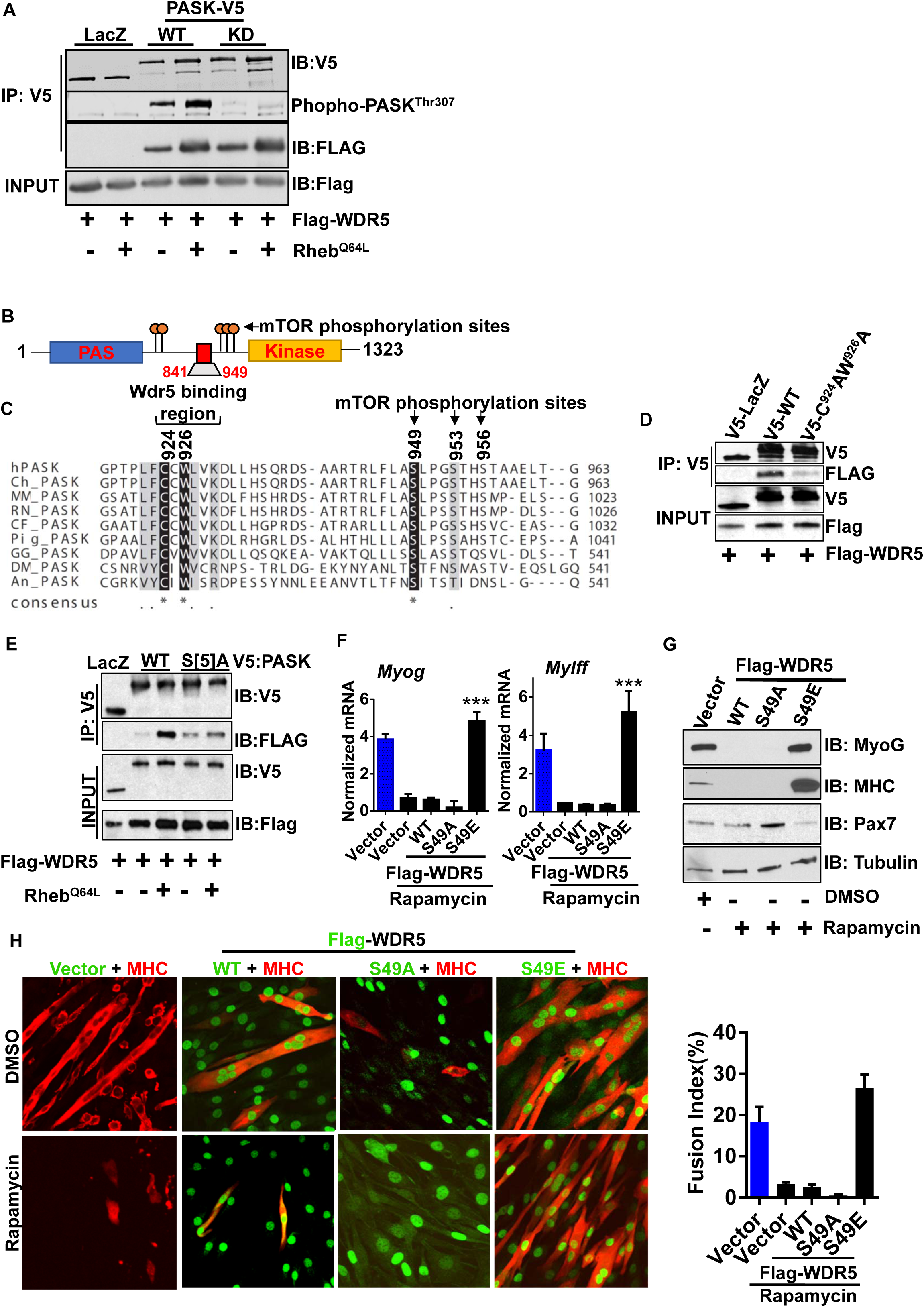
Wdr5 phosphorylation mediates mTORC1‐dependent myogenesis. **[A]** V5-LacZ as control or WT or KD PASK were co-expressed with Flag-tagged WT-Wdr5 in the presence or absence of Rheb^Q64L^. 24 hr after transfection, V5-tagged proteins were immunoprecipitated and abundance of Flag-Wdr5 was determined by western blotting. Activation of PASK by Rheb^Q64L^ was measured by western blotting of the immunoprecipitates with anti-AKT substrate antibody. **[B]** Domain arrangement of human PASK indicating relative positions of mTORC1-dependent phosphorylation sites and the Wdr5-interacting region on PASK. **[C]** Alignment of a region of PASK encompassing Wdr5 binding region and mTOR phosphorylation sites from different species. Conserved residues are marked by black boxes. **[D]** V5-tagged LacZ or WT or C^924^A/W^926^A PASK were co-expressed with Flag-Wdr5 in HEK293T cells. The association between Wdr5 and various PASK proteins was measured by probing immunoprecipitates using the indicated antibodies. **[E]** V5-tagged LacZ or WT or S[5]A (S^640^, T^642^, S^949^, S^953^, S^956^A) mutant PASK were co-expressed with Flag-Wdr5 in the presence or absence of Rheb^Q64L^. Western blotting to detect relative enrichment of Flag-Wdr5 was performed as in [A]. **[F]** C2C12 myoblasts were infected with retroviruses expressing control or WT, S^49^A or S^49^E Wdr5. 24 hr after infection, cells were treated with DMSO or 40 nM rapamycin for 24 hr followed by induction of differentiation by 100 nM insulin in the presence or absence of rapamycin as indicated. Normalized levels of mRNA for *Myog* and *Mylff* (Mhc) were determined using qRT-PCR with 18s rRNA used as normalizer. Blue bars indicate the extent of normal myogenesis in the absence of rapamycin inhibition for comparison. **[G]** Western blot analysis from an experiment as in [F]. **[H-I]** Immunofluorescence microscopic examination of myogenesis of WT and the indicated Wdr5 mutants in DMSO or rapamycin. Fusion index was calculated as described in methods.

**Figure 7.**
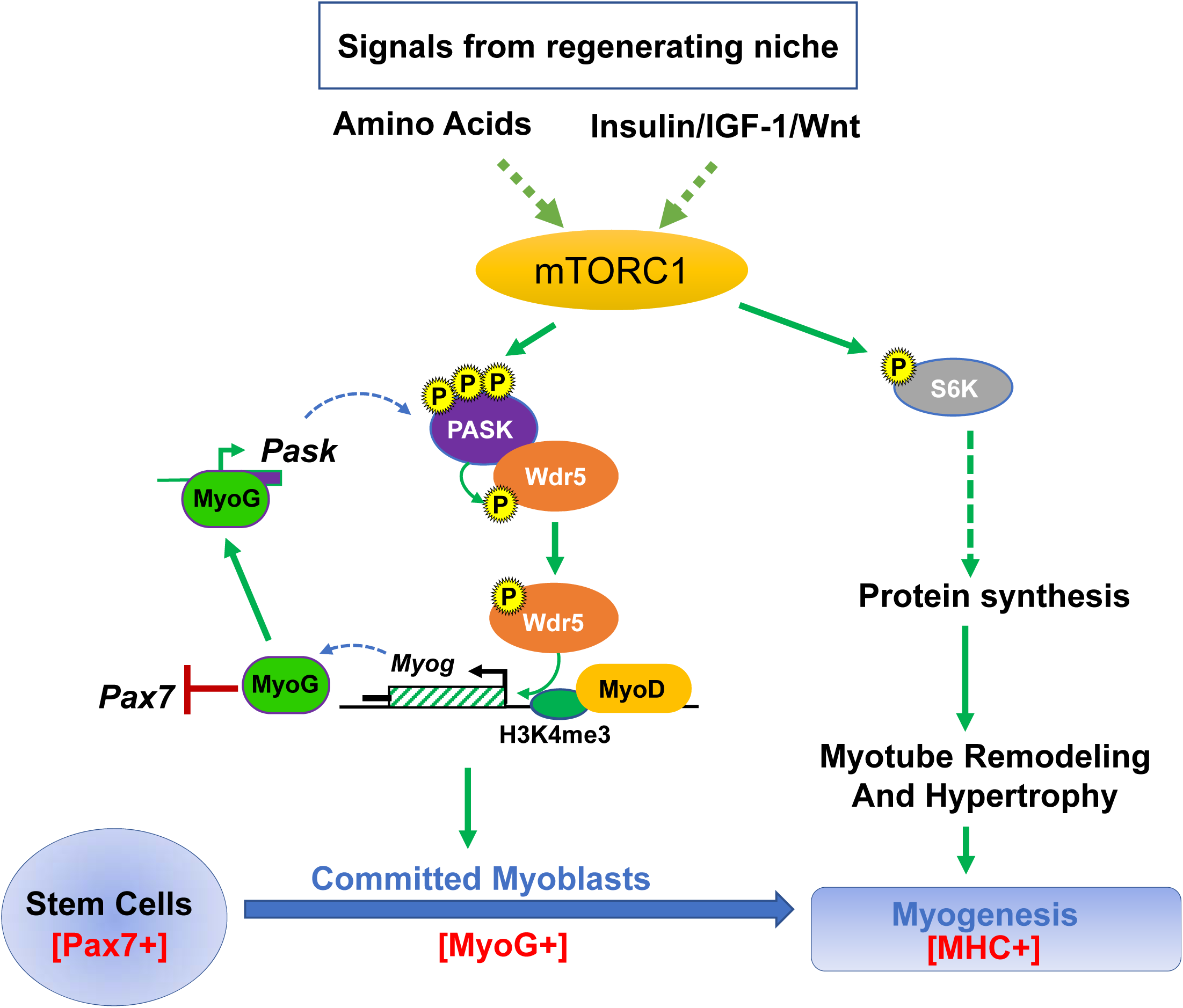
The role of mTORC1-PASK-WDR5 in myogenic signaling. mTORC1 controls both the onset (establishment of commitment) and remodeling stages of myogenesis. Our findings show that PASK is a key missing downstream effector of mTORC1 function in MuSCs that is required for the transition from Pax7+ stem cells to MyoG+ committed progenitors. Once the myogenesis program is induced, S6K1-driven translational upregulation results in myotube maturation and metabolic adaptation necessary for muscle function.

## Discussion

mTORC1 integrates multiple signals from the regenerating niche, including nutrients as well as hormones such as insulin, IGF-1 or Wnt, and is required for myogenesis and muscle repair. In this study, we show that PASK is a substrate of mTORC1 downstream of these same niche signals, particularly insulin and nutrients. mTORC1-dependent phosphorylation and activation of PASK stimulate its association with and phosphorylation of Wdr5. This Wdr5 phosphorylation triggers epigenetic changes at the *Myog* promoter that activate *Myog* transcription and, thereby, establish the commitment to myogenic differentiation (Figure *7*). In addition, MyoG also occupies and stimulates the *PASK* promoter to establish a positive feedback loop supporting the irreversible commitment to differentiate (Kikani et al., 2016). mTORC1 activation of p70S6K simultaneously activates the protein synthesis that is required for rapid myotube hypertrophy, resulting in the culmination of the myogenesis program. Thus, mTORC1 coordinately enables both aspects of myogenesis via activation of distinct protein kinase signaling pathways. mTORC1, PASK, and Wdr5 are widely expressed in stem cells. As PASK is required for differentiation of multiple stem cell lineages (Kikani et al., 2016), this model could represent a common mechanism by which nutrient and hormonal signaling could establish the commitment to differentiate via mTORC1 signaling. This might be particularly relevant for cell types that are highly metabolically active, like muscle cells and adipocytes (Martin et al., 2015).

The role of the mTOR protein kinase in the regulation of myogenesis appears to be multi-dimensional (Ge and Chen, 2012). mTOR has been shown to regulate the myogenesis program using two different mechanisms, only one of which depends upon its catalytic activity (Erbay and Chen, 2001; Ge and Chen, 2012). Moreover, mTOR regulates not only the early stages of myogenesis but also controls the remodeling of myotubes after differentiation (Risson et al., 2009; Zhang et al., 2015). Despite considerable interest, it remains unclear how mTOR signals to establish early steps of myogenic commitment in muscle stem cells. Multiple genetic studies and our data in Figure 4E suggest the requirement of the Raptor-containing mTOR complex 1 (mTORC1), but not the Rictor-containing mTORC2 to mediate the early steps of myogenic commitment (Bentzinger et al., 2008; Hung et al., 2014; Rodgers et al., 2014; Wan et al., 2006; Wu et al., 2015). One of the major substrates of mTORC1, p70S6K1 is dispensable for the early steps of the myogenesis program (Figure 4E and (Ge and Chen, 2012; Ohanna et al., 2005)). S6K1 or 2, however, play important roles in myotube remodeling. Our results suggest that PASK may be the missing downstream effector of mTORC1 signaling that initiates myogenesis. mTORC1 phosphorylates PASK within two distinct regions: the N-terminal residues S^640^ and T^642^ and the C-terminal residues S^949^, S^953^ and S^956^. These residues are highly conserved, suggesting the possible conservation of mTORC1-PASK signaling across metazoan species. Interestingly, similar to what is observed for several other mTORC1 substrates, the two regions of PASK sites have variable rapamycin sensitivity. Consistent with being more robust sites, phosphorylation at S^949^, S^953^ and S^956^ is rapamycin resistant. These sites are adjacent to the Wdr5 binding motif, and we showed that phosphorylation augments PASK interaction with Wdr5, and presumably activation of myogenic gene expression. It remains unclear what role phosphorylation of the N-terminal residues plays in PASK function, but they are near the conserved PAS domain, which binds and regulates the kinase domain of PASK. It is possible that mTORC1 signaling may impact the intramolecular interaction between the PAS and kinase domains. Thus, mTORC1 might function to integrate nutrient and hormonal signals to increase PASK activity and Wdr5 phosphorylation to coordinate myogenesis.

PASK expression was induced several-fold during regenerative myogenesis as early as day 3 post-injury. We and others have observed that this increase in *PASK* expression originates from proliferating MuSCs (Kikani et al., 2016; Liu et al., 2013) and is coincident with the induction of *Myog* expression (Rodgers et al., 2014). Our data presented here suggest that mTORC1-mediated post-translational activation of PASK might provide another layer of control over the critical decision of commitment to differentiate. We propose that such precise signaling control of the epigenetic network is required to regulate myogenesis in accordance with the environmental status. As such, multiple pharmacological and genetic approaches have established that hyper-activation of PI3K/mTORC1 signaling leads to MuSC exhaustion and exacerbates age-associated muscle wasting, presumably due to premature myogenesis and loss of self-renewal potential. Similarly, we show that over-expression of WT PASK (Figure 5A-D, S1B) or Wdr5^S49E^ is sufficient to activate the myogenesis program even in the absence of differentiation signaling. Thus, PASK activity and expression are both controlled to prevent precocious differentiation of MuSCs. By integrating nutrient information with epigenetic changes via Wdr5 phosphorylation, mTORC1-PASK signaling can precisely and appropriately control the myogenesis program.

The discoveries described herein enable a more thorough understanding of the mechanisms underlying the coordination of stem cell fate decisions with extracellular signaling through the mTORC1-PASK-Wdr5 pathway. To this end, our results establish a functional link between mTORC1 signaling and MLL and other histone-modifying complexes of which Wdr5 is a component. mTORC1 and MLL are two critical protein complexes within stem cells that are intimately involved in various aspects of stem cell functions (Aziz et al., 2010; Bulut-Karslioglu et al., 2016; Castilho et al., 2009; McMahon et al., 2007; Yilmaz et al., 2012; Zhang et al., 2016). The identification of PASK-Wdr5 as a mediator of mTORC1 effects on transcriptional programming set the stage for a deeper understanding of the possible cross-talk between these two complexes and how that impacts stem cell functions.

## Materials and Methods

### Generation of PASK transgenic mice

PASK tagged with V5 at the C-terminus was subcloned into the pBigT plasmid (a kind gift from Mario Capecchi) using the NheI and NotI sites (forward primer 5’-CTCACAGCTAGCGCCGCCACCATGGAGGACGGGGGCTTAACAGCC-3’, reverse primer 5’-GGAAGCGGCCGCTCACGTAGAATCGAGACCGAGGAG-3’). This generated a plasmid with V5-PASK downstream of *loxP-STOP-loxP.* The resulting construct was digested with PacI and AscI and the fragment was inserted into the pROSA-1PA plasmid (a kind gift from Mario Capecchi). The final construct was linearized with SalI and the digestion product was purified by phenol-chloroform extraction. Electroporation of C57Bl/6 ES cells and selection of ES clones was performed by the Transgenic and Gene Targeting Mouse Core at the University of Utah. ES cells were screened for successful targeting of V5-PASK by PCR (forward primer 5’-CCTAAAGAAGAGGCTGTGCTTTGG-3’, reverse primer 5’CATCAAGGAAACCCTGGACTACTG’), and by infecting cells with MSCV-Cre (a kind gift from Don Ayer), and analyzing cell lysates by Western blotting with anti-V5 and anti-PASK antibodies. Two correctly targeted clones were injected into blastocysts and generated germline chimeric mice. Genotyping was performed by PCR using DNA isolated from ear biopsies. Primer 1 (5’-GGAGGGGAGTGTTGCAATACCTTT-3’ and primer 2 (5’-AGCTGGGGCTCGATCCTCTAGTTG-3’) were used to detect wild-type and transgenic alleles respectively. The PCR program consisted of 2 min at 95°C, then 30 cycles of 30 s at 95°C, 30 s at 58.5°C, 1 min at 72°C and 5 min at 72°C. Mice were maintained on a C57Bl/6 background. ROSA26^PASKFLS/^+ mice were crossed to C57Bl/6 transgenic mice expressing Cre recombinase under a CMV promoter (a kind gift from the Transgenic and Gene Targeting Mouse Core at the University of Utah) to generate experimental animals (ROSA26^hPASK-V5^).

### Cell lines

HEK293E, HEK293T, and C2C12 myoblasts were obtained from ATCC. Parental *Tsc2*^−/−^ and h*Tsc2*-complemented MEFs were a gift from Dr. Brendan Manning. These cells were all cultured in DMEM supplemented with 10% Fetal Bovine Serum and 1% Penicillin and Streptomycin. For amino acid stimulation, custom DMEM was ordered from Invitrogen to lack L-leucine or L-leucine and L-arginine. For all nutrient and insulin stimulation experiments, either C2C12, HEK293E or MEFs were used as these cells respond to insulin. HEK293T cells were used for an experiment involving Rheb^Q64L^ overexpression. All cells were routinely checked for the presence of mycoplasma and were karyotyped by ATCC. All cells were kept at 37°C with 5% CO_2_.

### Isolation of primary myoblasts

MuSCs were isolated from 10-12 week old WT and *PASK*^−/−^ littermates according to published protocols (Danoviz and Yablonka-Reuveni, 2012). Briefly TA muscles from hind-limbs of WT or *PASK*^−/−^ mice were isolated, minced in DMEM and enzymatically digested with 0.1% Pronase for 1 hr. After repeated trituration, the cell suspension was filtered through a 40 μM filter. Cells were plated on Matrigel-precoated plates and allowed to grow for four days. The differentiation of these MuSC-derived myoblasts was stimulated by the addition of 100 nM insulin in serum-free DMEM.

### Insulin, amino acids and glucose stimulation

For insulin stimulation to measure PASK activity using HEK293E or MEFs, cells were serum-starved overnight (12 hr) in DMEM without FBS. Inhibitors such as rapamycin were also added during serum starvation as indicated. 100nM insulin or vehicle control was added directly to the culture media. Cells were incubated at 37°C for the times indicated in each experiment. For amino acid stimulation, cells were serum and amino acids starved overnight. Following, a freshly prepared solution of L-leucine was added directly to the culture medium at a 0.8mM final concentration. Similarly, for glucose stimulation, cells were cultured overnight in low glucose media (1g/L) followed by glucose free DMEM for 1hr. Cells were then stimulated with a final concentration of 25mM D-glucose.

### Cell lysis and immunoprecipitation

For immunoprecipitations, cells were lysed in a native lysis buffer containing 40mM HEPES pH 7.4, 150mM NaCl, 2mM KCl, 1mM EDTA, 1mM EGTA, 100mM Sodium pyrophosphate, 10mM β-glycerophosphate and protease and phosphatase inhibitor cocktails. Lysates were incubated on ice for 15 min followed by high-speed centrifugation to pellet insoluble debris. The soluble fraction was subjected to immunoprecipitation using the antibodies as indicated.

### In vitro kinase assay

The *in vitro* kinase assays were performed essentially as described previously (Kikani et al., 2010; Kikani et al., 2016). Briefly, endogenous or over-expressed PASK was purified from cells using anti-PASK antibody or anti-V5 antibody using native cell lysis buffer as described above. The kinase reaction was performed by washing immunoprecipitated PASK with Kinase buffer without ATP (20mM HEPES 7.4, 10mM MgCl_2_, 50 μM ATP, 1mM DTT). The kinase reaction was initiated by adding 100ng of purified Wdr5 or Ugp1 as substrate and 1 μCi/reaction of [γ-^32^P]ATP (PerkinElmer Life Sciences). The reaction was terminated after 10 min by adding denaturing SDS sample buffer. The proteins were separated by SDS-PAGE, transferred to a nitrocellulose membrane and visualized by autoradiography. Western blot analysis using anti-PASK or anti-V5 antibody was performed on the same membrane to determine loading for normalization purposes.

### Metabolic in vivo labeling

Metabolic *in vivo* labeling was performed in cells expressing various PASK plasmids with or without constitutively active Rheb (Rheb^Q64L^). 24 h after transfection, cells were washed twice with phosphate-free DMEM (Invitrogen) followed by incubation with 1.0 mCi of ^32^P. Cells were washed with phosphate-free DMEM to remove unincorporated ^32^P, lysed using lysis buffer, and immunoprecipitated as described above. Immunocomplexes were washed with buffer (20 mM Na_2_HPO_4_, 0.5% Triton X-100, 0.1% SDS, 0.02% NaN3) containing high salt (1M NaCl and 0.1% BSA) followed by low salt (150 mm NaCl) in the same buffer. Immuno-precipitated PASK was released by SDS-PAGE sample loading buffer, separated by SDS-PAGE, followed by transfer to nitrocellulose membrane and autoradiographic imaging.

### Immunofluorescence microscopy

Primary myoblasts isolated from *Rosa26^hPASK-V5^* or WT mice or C2C12 myoblasts growing on coverslips were fixed with 4% Paraformaldehyde and permeabilized with 0.2% Triton-X100. Following 1 hr of blocking with 10% normal goat serum, the indicated primary antibodies were added for overnight incubation at 4°C. Following three washes with ice cold PBS, cells were incubated with anti-mouse Alexafluor 568 (for MyoG and MHC) or anti-rabbit Alexafluor 488 (for Flag or V5-PASK or Flag-Wdr5) secondary antibodies for 1 hr in the dark at room temperature. The coverslips were mounted using Prolong-Antifade mounting medium containing DAPI. The fusion index was used as a measure of differentiation and was calculated as the percent of total nuclei in MHC^+^ cells. For quantification of microscopic images, at least 100 cells were counted from three separate experiments in a blinded manner. Statistical significance was calculated using Student's t-test with p<0.05 set as the significance level.

## Supplementary Figure Legend

**Figure S1.**
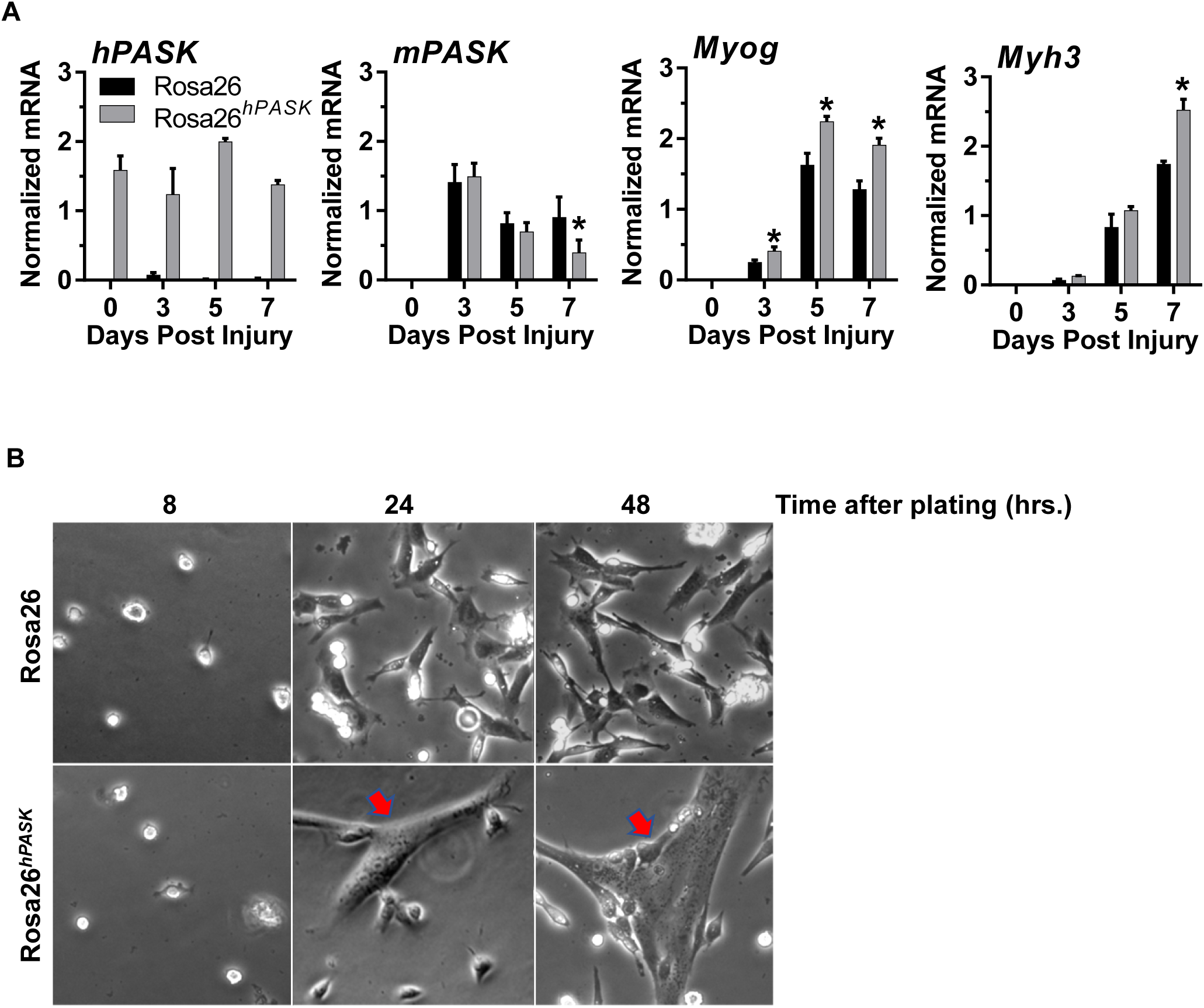
**(A)** Enhanced expression of myogenic genes during skeletal muscle regeneration of *Rosa26^hPASK-V5^* mice. Mice from the indicated genotypes were injured by intramuscular injection of 25μ! of 1.2% BaCh into the left TA muscles of 12 weeks old mice. Uninjured mice were used as Day 0 control. Mice were euthanized at days 3, 5 or 7 post-injury and muscles were isolated to prepare mRNA extracts. The relative abundance in the indicated transcripts was quantified by qRT-PCR using specific oligonucleotide primer sets. At least three mice were used for each time point. Error bars represent ±S.D. \**P*<0.05. **(B)** MuSCs were isolated from *Rosa26^hPASK-V5^* mice or littermate controls. Cells were allowed to grow in growth media and micrographs of cells were taken at the indicated time points. Red arrows indicated myotubes.

